# PhIP-Seq uncovers novel autoantibodies and unique endotypes in interstitial lung disease

**DOI:** 10.1101/2023.04.24.538091

**Authors:** Vaibhav Upadhyay, Young me Yoon, Sara E. Vazquez, Tania E. Velez, Kirk D. Jones, Cathryn T. Lee, Christopher S. Law, Paul J. Wolters, Seoyeon Lee, Monica M. Yang, Erica Farrand, Imre Noth, Mary E. Strek, Mark Anderson, Joseph DeRisi, Anne I. Sperling, Anthony K. Shum

## Abstract

Interstitial lung diseases (ILDs) are a heterogeneous group of disorders that can develop in patients with connective tissue diseases (CTD). Establishing autoimmunity in ILD impacts prognosis and treatment. ILD patients are screened for autoimmunity by assaying for anti-nuclear autoantibodies, rheumatoid factors and other non-specific tests. However, this approach has not been rigorously validated and may miss autoimmunity that manifests as autoantibodies to tissue antigens not previously defined in ILD. Here, we use Phage Immunoprecipitation-Sequencing (PhIP-Seq) to conduct a large, multi-center unbiased autoantibody discovery screen of ILD patients and controls. PhIP-Seq identified 17 novel autoreactive targets, and machine learning classifiers derived from these targets discriminated ILD serum from controls. Among these 17 candidates, we validated Cadherin Related Family Member 5 (CDHR5) as an autoantigen and found CDHR5 autoantibodies in patients with rheumatologic disorders and importantly, subjects not previously diagnosed with autoimmunity. Lung tissue of CDHR5 autoreactive patients showed transcriptional profiles consistent with activation of NFκB signaling and upregulation of chitotriosidase (CHIT1), a molecular pathway linked to fibrosis. Our study shows PhIP-Seq uncovers novel autoantibodies in ILD patients not revealed by standard clinical tests. Furthermore, CDHR5 autoantibodies may define a novel molecular endotype of ILD characterized by inflammation and fibrosis.

## INTRODUCTION

Interstitial lung diseases are a heterogeneous group of disorders with well-defined disease associations and genetic underpinnings (Liu et al., 2023; Wijsenbeek et al., 2022; Hyldgaard et al., 2017; Raghu et al., 2018). Connective tissue disorders (also referred to as systemic rheumatic disorders) such as rheumatoid arthritis (RA) and scleroderma are known causes of ILD (Graney and Fischer, 2019). In patients with a CTD, ILD often has the greatest impact on morbidity and mortality (Hyldgaard et al., 2017; Morisset et al., 2017). Detecting whether autoimmunity is present in an ILD patient has important implications for treatment and prognosis, although identifying autoimmunity can be difficult in patients that do not meet the definition of a known CTD (Graney and Fischer, 2019).

Expert guidelines recommend that every ILD patient should be screened for a CTD by assessing for serologic markers (Fischer et al., 2015; Graney and Fischer, 2019; Raghu et al., 2018). However, there is no clear consensus as to which tests should be performed (Raghu et al., 2018). Most of the laboratory studies that are recommended (anti-nuclear antibodies, C-reactive protein, erythrocyte sedimentation rate, rheumatoid factor) lack specificity and their use for the detection of autoimmunity in the context of ILD has not been rigorously studied (Raghu et al., 2018; Graney and Fischer, 2019).

Autoantibodies utilized to diagnose systemic rheumatic disorders may be insufficient to screen for autoimmunity in ILD patients. Pulmonologists and rheumatologists have recognized for years that autoimmune-associated ILD may in fact be a unique disease (Vij et al., 2011; Kinder et al., 2007). Importantly, ILD is not considered a criterion for nearly all the defined CTDs in which it manifests. There is a well-described subset of ILD patients with clinical features of autoimmunity that do not meet criterion for a defined CTD (Fischer et al., 2015). Thus, it is possible that for some ILD patients there are autoantibodies unique to ILD that are missed by standard lab tests used to assess for an underlying rheumatologic condition.

Furthermore, detecting autoantibodies outside the context of a defined CTD is increasingly recognized as relevant (Salvator et al., 2022; Sinnberg et al., 2023). Recent data shows that such autoantibodies can significantly impact patient outcomes, even in clinical settings in which autoimmune mechanisms are not the primary driver of disease. In COVID-19, autoantibodies to type I IFNs predispose individuals to severe disease, including significant lung damage (Bastard et al., 2020, 2021; van der Wijst et al., 2021). Thus, it is possible that novel tissue autoantibodies in ILD patients, including in those without a defined CTD, may have an important role in disease pathogenesis.

Phage Immunoprecipitation Sequencing (PhIP-Seq) is an emerging technology enabling massively parallel profiling of autoreactive antibodies in patient serum by pairing programmable phage-display libraries with next generation sequencing (Mandel-Brehm et al., 2019; Vazquez et al., 2020, 2022; Larman et al., 2011). PhIP-Seq facilitates the profiling of serologically reactive peptide epitopes tiling all open-reading frames of the human genome. PhIP-Seq has been applied to define novel autoantibodies in established diseases and has provided new insight into pathogenic mechanisms and new autoimmune syndromes (Mandel-Brehm et al., 2019, 2022). In this study, we hypothesized that PhIP-Seq could discover novel autoantibodies in ILD to detect autoimmunity not revealed by standard tests and define new endotypes of disease.

## RESULTS AND DISCUSSION

### Study population

To discover autoantibodies linked to ILD, we paired the unbiased profiling strategy of PhIP-Seq with a large collection of patient samples enriched for commonly used ILD classifications. We selected patients from two academic medical centers with established ILD programs and experience in the multidisciplinary diagnosis of ILD. Because our goal was to discover novel autoantibodies irrespective of whether a defined CTD had been established in a patient, we started with a heterogenous group of subjects. To control for variability in diagnostic agreement between the multidisciplinary diagnosis of the institutions, we specifically looked for autoantibodies shared between centers (see *Methods*).

We screened 398 ILD subjects and 138 individuals whose serum was obtained from blood banks that formed our reference cohort for autoantigen candidate selection (Vazquez et al., 2022). We included an additional group of patients with RA without known ILD (n=15) and subjects who were relatives of ILD subjects without known ILD (n=16) or organ donors from Center 2 (n=15) whose lungs were not used for transplantation; because these latter samples all had some clear biological feature that makes them distinct from the blood donors who were used as the reference population, we include them as a separate group of subjects recruited to Center 2 but who do not have ILD (Center 2 Non-ILD). Including technical replicates, we completed PhIP-Seq on 704 total samples from 582 subjects (**Table 1**). Screened subjects represented a broad array of diagnostic categories (**Table 1**), which reflected major diagnoses commonly referred to and treated at the ILD centers included in this study. We studied similar numbers of male (48.1%) and female (49.8%) participants. While the majority of ILD subjects were White (69.1%), our screen included representation from multiple distinct demographic groups assessed by self-identified race/ethnicity categories (**Table 1**).

### PhIP-Seq identifies novel autoreactive peptide targets in ILD subjects

From the PhIP-Seq screen, we selected autoantibody candidates by utilizing stringent cutoffs for autoreactive peptide targets with high levels of reactivity across ILD programs (see *Methods*), reasoning these would be most likely to validate in orthogonal assays and be clinically meaningful. Using this approach, we identified 17 candidate autoantigens shared between ILD programs (**Figure 1A, Supplementary Figure 1A**). Nine of these 17 had little to no reactivity in the Center 2 Non-ILD group (**Figure 1A**) with the remaining 8 having reactivity in this group similar to that seen in the ILD subjects (**Supplementary Figure 1A**). We compared our data to autoantibodies previously reported in the ILD literature (**Supplementary Figure 1B)**. While PhIP-Seq was able to detect a subset of these, none of the historical autoantibodies met the same benchmarks we instituted in this study. Notably, of the 17 autoantigen candidates that met our screening criteria, to our knowledge none have been previously reported in studies of systemic rheumatic disorders or other lung diseases.

**Figure 1:**
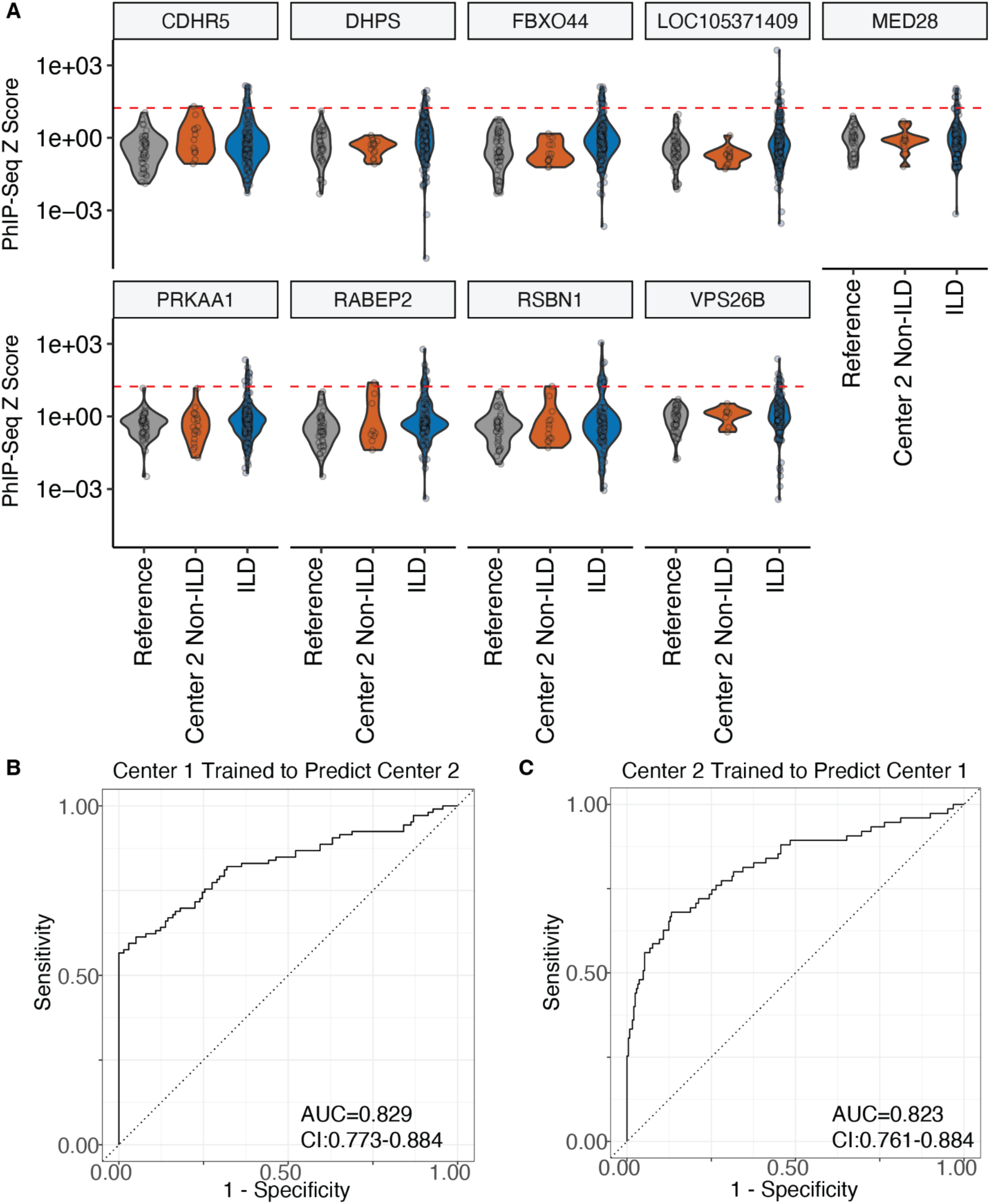
PhIP-Seq identifies novel candidate autoantigens with predictive capacity for ILD. **(A)** Candidate autoantibodies were selected using Z score cutoffs (red dashed line, Z score cutoff = 17) from reference samples and subjects from two separate ILD centers. Additional subjects were recruited from ILD Center 2 including a mixture of subjects not known to have established ILD (n=138 reference, n=398 participants with ILD, n=46 participants in the group Center 2 Non-ILD). **(B-C)** A random forest classifier was trained on PhIP-Seq data derived from all screened subjects and trained either on **(B)** Center 1 to predict Center 2 or **(C)** vice versa to distinguish between subjects with and without ILD. Patients with known RA without known ILD were excluded from **(B-C)** given potential overlap of autoreactivity with patients with RA and ILD. Area under the curve (AUC) and 95% confidence interval (CI) are annotated on the graphs. **(B-C)** n=169 control subjects and n=398 subjects with known ILD.

A close examination of our candidates revealed that many but not all the autoantigens targeted are derived from proteins expressed within lung tissue, and the proteins have a variety of molecular functions or localize to a variety of cellular compartments (Joshi et al., 2020; DePianto et al., 2015; Lu et al., 2014; Bodempudi et al., 2014). Overall, the candidate autoantigens we discovered were diverse and without easily discernible relationships to one another, aside from their PhIP-Seq derived autoreactivity in ILD patients. The number of patients demonstrating autoreactivity reflected approximately 2.5% of the total cohort and included patients from all the groups represented in the screen, including those with and without defined autoimmune syndromes (**Figure 1A** and **Supplementary Figures 1-2**). Because the patients and autoantibodies were heterogeneous, we performed a computational analysis of the data using machine learning to uncover complex associations not otherwise apparent after initial review.

### PhIP-Seq derived candidate autoantigens discriminate between ILD subjects and controls

Previously, a logistic regression derived machine learning algorithm has been successfully used to discriminate cases from controls using PhIP-Seq input data with respect to a hereditary form of autoimmunity (e.g. APS1 as described in (Vazquez et al., 2022)). Here, we trained a random forest classifier to ask how PhIP-Seq selected candidate autoantigens would perform as a serum based test to predict the presence of ILD. We developed our classifier using PhIP-Seq data for the 17 candidates where the classifier was trained on the data from one center and tested on the other, or vice versa (**Figure 1B-C**). Control samples included the reference group (n=138 subjects) and samples from the group designated Center 2 Non-ILD without RA (n=31 subjects). We utilized samples from all patients with ILD (n=398 subjects). Using this approach, we found PhIP-Seq derived autoantibodies maintained the capacity to distinguish between ILD and control samples even when the center of origin was distinct (AUC=0.829, CI: 0.773–0.884 and AUC=0.823, CI: 0.761–0.884) (**Figure 1B-C**). These findings indicate that in the future clinicians without access to a multidisciplinary diagnosis may be able to verify the presence of ILD through a blood test derived from the PhIP-Seq autoantibodies. Taken together, PhIP-Seq uncovered a panel of candidate autoantigens that are both novel and unique to ILD patients.

### A subset of ILD subjects have autoantibodies to an immunodominant epitope of CDHR5

We next examined whether any of the autoantibodies could define a specific subset of ILD subjects. We focused on the most unique autoantibodies, reasoning that a highly specific autoantibody that arises in the context of a restricted immune response was more likely to define a disease subtype. First, we built a network diagram of interactions to indicate how frequently an autoantigen candidate was found in association with other candidates across samples (**Figure 2A**). All the autoantigen candidates in our panel occurred in association with at least one other and none were found in isolation. However, when we utilized the specific peptide epitopes for our analysis (which are filtered to Z score cutoffs of 50) we found Cadherin Related Family Member 5 (CDHR5) had the most unique profile (**Figure 2B-C**). All but one of the 11 unique CDHR5 peptides comprised overlapping amino acid sequences that mapped to amino acids 155-271 of the protein (**Supplementary Table 1**). Further analysis of the predicted CDHR5 secondary structure revealed that a putative beta-pleated sheet predicted to be an internal domain of the protein is likely the immunodominant epitope targeted by 10 out of 11 autoantibodies (**Figure 2D**).

**Figure 2:**
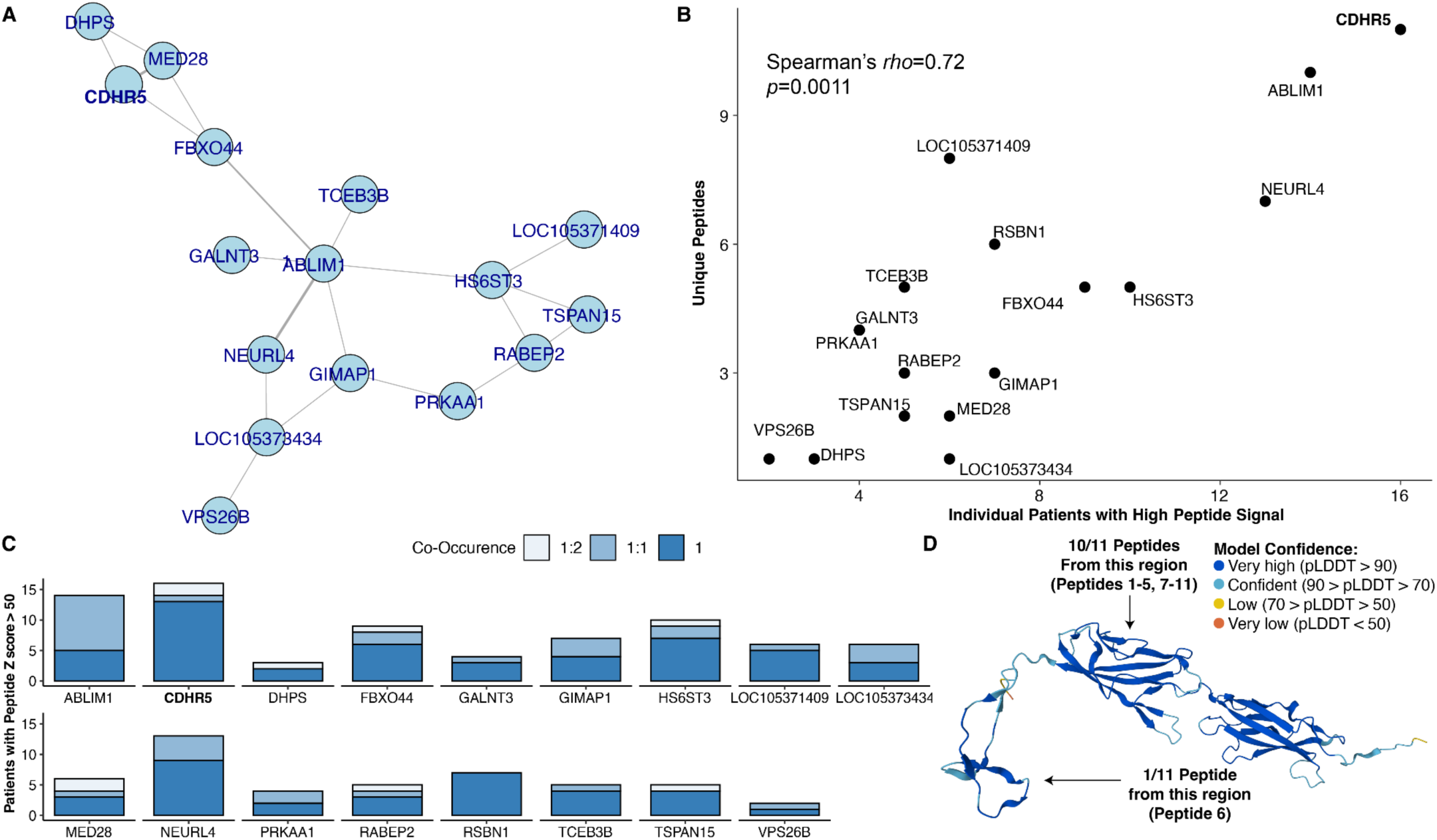
CDHR5 is a unique candidate autoantigen in a subset of ILD subjects. **(A)** Network of interactions for data transformed into peptides above a Z score cutoff of 50 per individual patient. Each edge indicates co-occurence of given candidates within a screened subject. Each node represents a gene for which peptides were detected. **(B)** Number of unique peptides for a given autoantigen is plotted against the number of unique patients with peptide autoreactivity. A Spearman’s correlation coefficient and *p*-value is annotated. **(C)** Co-occurence of both gene and peptide data by total number of positive subjects with darker intensity colors indicating uniqueness (1:2 indicates the designated autoantigen was found with two other autoantigen candidates in a given patient; 1:1 indicates indicates the designated autoantigen was found with one other autoantigen candidates in a given patient; 1 indicates the specified autoantigen was found in the absence of the remaining 16 candidate autoantigens). **(D)** Predicted structure of protein using AlphaFold with UniProt accession A0A7L0FKT8; per-residue confidence score (pLDDT) between 0 and 100 coloring indicated in legend inset. Peptide annotations as described in **Supplementary Table 1**.

Patients demonstrated varying levels of reactivity to CDHR5 based on PhIP-Seq Z scores (**Supplementary Figure 2**). For CDHR5, subjects with hypersensitivity pneumonitis (HP) had significantly higher Z scores than those with idiopathic pulmonary fibrosis (IPF) or other forms of ILD, and patients with CTD-ILD showed *p*-values that exhibited trends in the same direction (**Supplementary Figure 2**). None of the other candidate autoantibodies revealed any discriminatory behavior by diagnoses when analyzed similarly (**Supplementary Figure 2**, Kruskal-Wallis *p*>0.05). For some subjects with CDHR5 autoreactivity, these data suggest a clinical syndrome similar to CTD-ILD or HP may be identifiable (**Supplementary Figure 2** and **Table 2**). Taken together, CDHR5 appeared to be an autoantibody candidate with strong potential to define a novel subtype of ILD. We therefore sought to validate this further with an orthogonal experimental approach.

### CDHR5 autoantibodies are elevated in ILD subjects with and without known autoimmunity

To confirm the presence of CDHR5 autoantibodies in patient serum, we developed a CDHR5 radioligand binding assay (RLBA). Using RLBA, we screened all the serum from patients and controls and validated the presence of CDHR5 autoantibodies in 18 ILD subjects with this orthogonal approach (**Figure 3A**). CDHR5 autoantibody-positive subjects were characterized by a number of ILD categorizations (**Figure 3B and Table 2**). Half of the subjects (9/18) had prior evidence of autoimmunity and a defined systemic rheumatic disease including mixed connective tissue disease (MCTD, n=4), RA (n=2), anti-synthetase syndrome (n=1) and scleroderma (n=1). Importantly, CDHR5 autoantibodies were detected in 9 of 18 patients without known autoimmunity, including one subject who met criteria for idiopathic pneumonia with autoimmune features (IPAF) and two subjects with unclassifiable ILD. Other diagnoses included HP (n=3), IPF (n=2) and interstitial lung abnormality (ILA, n=1) (**Table 2**).

**Figure 3:**
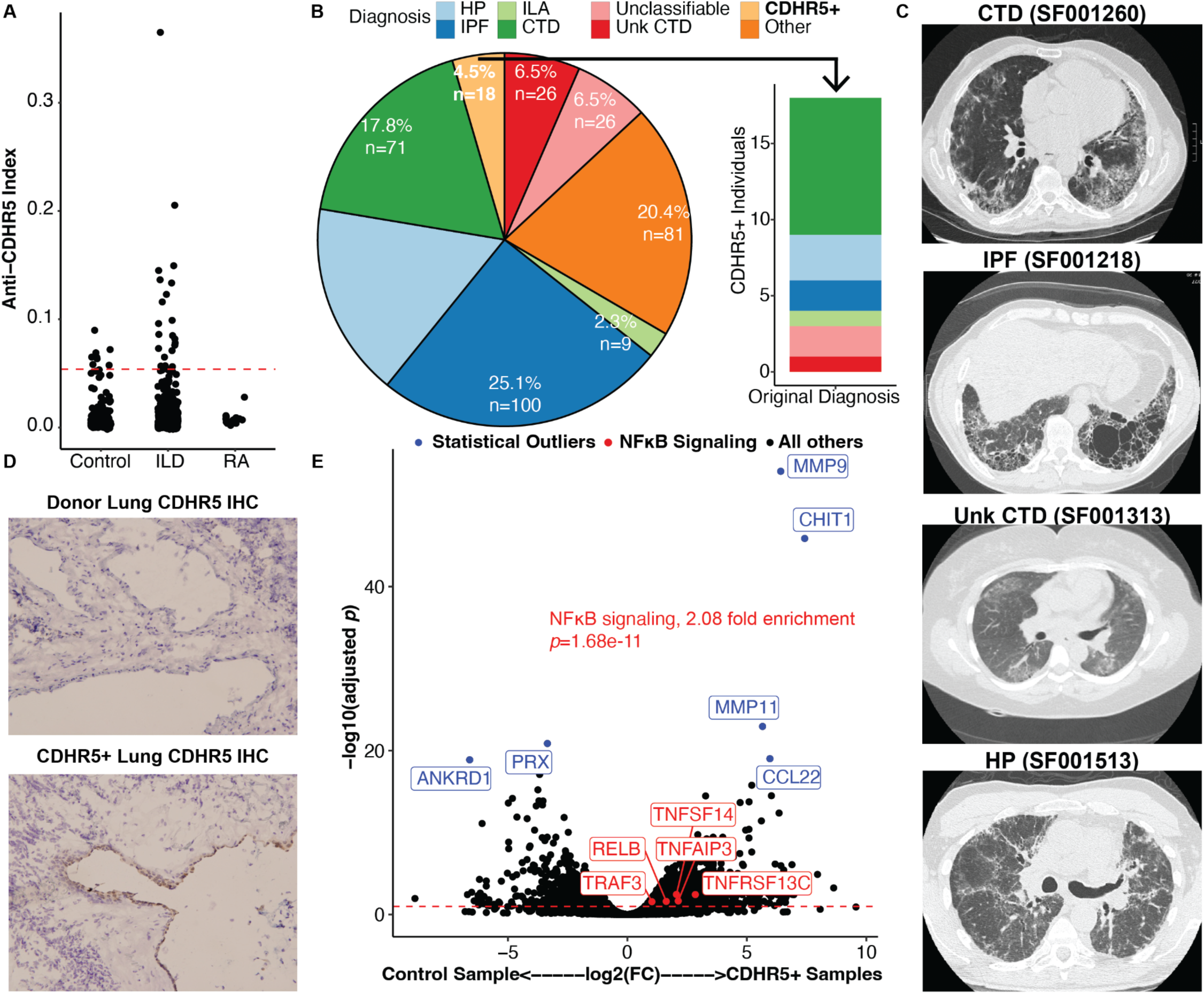
CDHR5 autoantibodies identify subjects with inflammation and fibrosis in the lung. **(A)** RLBA results for CDHR5 reactivity (Anti-CDHR5 Index on y-axis) separated by serum from indicated tested groups. Subjects are separated by having RA without known ILD (RA), all other non-ILD samples as Control, and all subjects with ILD in the ILD group (n=222 Control samples, n=454 ILD samples, n=15 samples for patients with RA without known ILD). **(B)** CDHR5 autoreactive subjects are indicated by the light orange wedge in the pie chart (CDHR5+, left) and the breakdown of these patients based on their original diagnoses is shown in the bar chart (right). **(C)** Representative images of CT scans from four different ILD diagnostic categories for the CDHR5+ subjects. **(D)** CDHR5 immunohistochemistry from unused donor lung and CDHR5 Ab+ subject lung tissue **(E)** Bulk RNA sequencing data for CDHR5 autoreactive subject lung tissue (n=2 subjects, SF001218 and SF001260) and age matched control tissue (n=3 subjects) analyzed via DeSeq2. Points represent individual RNA targets and the dashed line indicates significance (Benjamini Hochberg adjusted *p*<0.1). Significant outlier data points by *p*-value are highlighted in blue and pathway enrichment results for the most statistically enriched pathway, NFκB signaling, are annotated with a subset of molecular targets highlighted in red.

We performed detailed chart review of patients with CDHR5 autoantibodies and analyzed clinical, radiographic, and pathologic data (**Table 2** and **Figure 3C**). For all subjects classified as HP, a culprit antigen for inciting the disease was never identified and the diagnosis was not considered definitive for any patient (Raghu et al., 2020). Of note, 14 out of 18 subjects with autoreactivity to CDHR5 were either deceased or underwent lung transplantation at the time of writing this manuscript (**Table 2**). Ten of the deceased participants died or underwent lung transplant within approximately 5 years of study enrollment or first blood draw (**Table 2**). While CDHR5 autoreactive subjects with known CTD were uniformly given immunosuppressive therapy, subjects without a known CTD had more variable treatments, including antifibrotic medications or treatment of gastroesophageal reflux disease (**Table 2**).

### CDHR5 autoantibodies define a unique ILD endotype marked by lung inflammation and fibrosis

Although CDHR5 is not known to be expressed in respiratory tissue, we wanted to assess whether we could detect CDHR5 protein in the lungs of subjects with confirmed CDHR5 autoreactivity. We obtained lung tissue from a subject with CDHR5 autoantibodies and conducted immunohistochemistry using a commercially available antibody against CDHR5. CDHR5 staining was strongly detected in the respiratory bronchial epithelium the CDHR5 autoreactive subject (n = 1 out of 1 subjects), but notably not in the lungs of control subjects (n = 1 out of 3 subject with a low degree of CDHR5 autoreactivity) (**Figure 3D**). These data raise the possibility that CDHR5 autoantibodies circulating in a subset of ILD patients may contribute to lung disease pathogenesis by reacting to CDHR5 protein that has been upregulated in the bronchial epithelium.

We next performed bulk RNA sequencing of lung tissue from available two CDHR5 autoreactive subjects (n=2 subjects) and compared these results to data derived from age matched normal lung tissue (n=3 subjects) (Bhattacharya et al., 2021). We discovered 2,268 genes differentially expressed between the groups (Benjamini-Hochberg adjusted *p*<0.1, **Figure 3E**). In the two patients with CDHR5 autoantibodies, differential gene expression analysis of RNA transcripts from the most significantly upregulated genes revealed biological pathways linked to NFκB signaling including a number of transcripts that encode for receptor, ligand, and signaling mediators of the NFκB pathway (**Figure 3E**). Independent of the pathway analysis, the most significantly differentially expressed genes by adjusted *p*-value included chitotriosidase (CHIT1), which has been linked to fibrotic lung disease in recent studies (Chang et al., 2020; Van Dyken et al., 2017).

## DISCUSSION

In this study, we utilized PhIP-Seq to perform one of the largest unbiased autoantibody discovery screens in ILD. Because the diagnosis of ILD is challenging with potential for misclassification, we studied a heterogeneous cohort of patients, irrespective of whether patients had a confirmed (or suspected) systemic rheumatic disorder. We identified 17 novel candidate autoantigens associated with ILD, none of which have been previously described. The validated candidate, CDHR5, was found in patients with defined autoimmune disorders providing a direct link to conditions of impaired immunological tolerance in this patient population. We also discovered CDHR5 autoreactivity in patients without an established autoimmune disease. This raises the possibility that the autoantibodies can uncover occult autoimmunity not otherwise captured during a multidisciplinary ILD evaluation, including one performed at an established ILD program. Our data also suggest that standard serologic tests ordered during a routine ILD evaluation are insufficient to capture unique subtypes of autoimmune-associated ILD, particularly those that are distinct from established rheumatologic diseases.

Importantly, recent studies in COVID-19 (Bastard et al., 2020, 2021; van der Wijst et al., 2021) and other disease settings (Salvator et al., 2022; Sinnberg et al., 2023) have shown that autoantibodies can have a potent role in disease pathogenesis, even in the absence of a systemic rheumatic disease. Autoantibodies to type I interferons (Bastard et al., 2020, 2021; van der Wijst et al., 2021) or surfactant proteins (Sinnberg et al., 2023) can worsen lung damage either through their direct effect on anti-viral immune responses or by the physiological derangements they provoke in the lung. Although the precise mechanisms by which these types of autoantibodies arise are not well understood, these studies highlight the importance of autoantibody discovery in patients not typically suspected of harboring pathogenic autoantibodies. Although we have not yet shown that CDHR5 autoantibodies mediate fibrosis or inflammation, we validated it as a novel autoantigen and discovered autoreactivity to 16 other potential autoantigens in patients with ILD. It will be important to examine in future studies whether the autoantibodies we identified contribute to ILD even in patients for whom autoimmunity is not thought to be the primary driver of disease (e.g. IPF subjects).

The diagnosis and classification of ILD is challenging and the practice at specialized centers involves a multidisciplinary discussion of each patient integrating clinical, pathological, and radiographic information by experts (Travis et al., 2013). Because diagnostic agreement between multidisciplinary discussion of institutions can vary (Walsh et al., 2016), objective measures can minimize misclassification of ILD. Specific autoantibodies help fill this need by enabling clinicians to diagnose subtypes of ILD that provide insight into treatment and prognosis (e.g. MDA-5 autoantibody-positive dermatomyositis ILD (Sato et al., 2009; Nombel et al., 2021)) or mechanistic links to certain defects in immune tolerance (e.g BPIFB1 autoantibodies and central tolerance defects (Shum et al., 2013; Ferré et al., 2019)). The in-depth analysis of CDHR5 autoantibodies we performed in this study hints at the potential of PhIP-Seq to uncover a unique endotype of ILD that unifies seemingly disparate patients into a single group. In addition, our computational analysis using machine learning suggests that in the future the PhIP-Seq autoreactivity we identified might be employed by clinicians without access to an multidisciplinary discussion as a first step to confirm the presence of ILD. Establishing the PhIP-Seq panel as an objective, non-invasive tool in the work-up of ILD will require further investigation and validation.

While little is known about CDHR5, a prior gene ontology analysis revealed extensive association between CDHR5 and genes involved in transmembrane localization or intracellular transport (Bläsius et al., 2017), which suggests this protein may be exposed on the cell surface and lead to an antigen-specific immune response. Remarkably, 14 out of 18 of CDHR5 autoreactive subjects were either deceased or underwent lung transplantation at the time of writing of this manuscript (**Table 2**). Molecular analyses of patient tissue showed CDHR5 protein staining in a subset of respiratory bronchial epithelium and expression of RNA related to inflammation and fibrotic pathways. The staining in **Figure 3D** is non-specific though appears to show a membrane localization pattern, which if true would support the hypothesis that cell surface derived CDHR5 may be an antigen stimulating an autoantibody response; alternatively, future efforts securing non-fixed tissues could reveal CDHR5 to be an intracellular protein whose peptide fragments may be displayed as antigens through the major histocompatibility complex. We detected a mixed signature of both NFκB signaling and upregulation of CHIT1 and MMP9 and MMP11 transcripts in the lung tissue of two CDHR5 autoreactive subjects, suggesting peripheral CDHR5 autoreactivity may be a potential biomarker of subjects in whom there is both fibrosis and inflammation occurring within lung tissue. While inflammation is not the primary etiology of fibrosis in most subjects, inflammation can exacerbate fibrosis (Wynn, 2011; Wilson and Wynn, 2009). Multiple NFκB signaling targets and members of the tumor necrosis superfamily (TNF) were overexpressed within CDHR5 autoreactive subjects and may be amenable to experimental manipulation (e.g. TNFAIP3, TNFRSF13, TNFSF14, RELB) (Ma and Malynn, 2012). These could be key pathways around which future studies are designed to determine the degree to which NFκB signaling pathways impact fibrotic progression.

Our current study has limitations. PhIP-Seq identifies epitopes encoded in linear stretches of DNA and is not designed to capture post-translational modifications, which may have a role in autoimmune syndromes (Papini, 2009). In addition, we only had lung tissue available from two subjects for transcriptional characterization and immunohistochemistry, which limits the generalizability of the identified disease signatures to larger populations. Moreover, while we were able to successfully detect CDRH5 transcript in lung tissue, which in and of itself is a novel finding, we did not find a significant difference between control subjects and CDHR5 autoreactive subjects, possibly due to a lack of power given the low number of available biopsy samples. Finally, while we conducted detailed chart reviews, we were restricted in our ability to define additional autoantibodies that can be evaluated using clinical tests in CDHR5 autoreactive subjects because this information was not available in the electronic medical record and patient serum was limited.

Our work shows how an unbiased approach in patient selection and screening technology can identify novel autoantibodies that may uncover new endotypes of ILD outside the framework of established rheumatologic diseases. Future studies will be needed to determine whether the autoantibodies discovered here can help with improved identification, molecular understanding, and even treatment of patients with previously poorly understood subsets of ILD.

## ACKNOWLEDGEMENTS

Funding was provided by the National Institutes of Health (R01AI68299, R01AI137249 to A.K.S, K08HL165106 to V.U., 5T32AI007090-44 to Y.Y., and 5U19AI162310-02 to A.S.), the American Heart Association (T.V.), The Pulmonary Fibrosis Foundation (C.T.L.), The Nina Ireland Program for Lung Health at UCSF (P.W.), the UCSF Clinical Translational Sciences Institute (V.U.).

Conceptualization, V.U., A.K.S., A.S..; Data curation, V.U.; Formal Analysis, V.U.; Funding acquisition, A.K.S., V.U..; Investigation, V.U., Y.Y., S.V., T.V., K.D.J, C.T.L, C.S.L, S.L., M.Y.; Methodology, V.U., S.E.V., Y.Y., J.D.; Project administration, A.K.S, A.S, M.A., J.D. ; Resources, A.K.S, A.S., J.D, ; Software, S.E.V, J.D.; Supervision, P.J.W., E.F., A.K.S., I.N., M.E.S., M.A., J.D., A.S. A.K.S..; Validation, Y.Y.; Visualization, V.U.; Writing – original draft, V.U.; Writing – review & editing, A.K.S.

P.W. received grants from Roche, Sanofi, Pliant, Boehringer Ingelheim and personal fees from; Sanofi, Boehringer Ingelheim, and Blade Therapeutics. M.E.S. received grants and editorial assistance from Boehringer Ingelheim and personal fees from Fibrogen. M.S.A. owns stock in Medtronic and Merck and has consulted for Sana and Imcyse. J.D. is a paid scientific consultant for The Public Health Company, Inc., Allen & Co., and Delve Bio. There is no direct overlap between the current study and these consulting duties. The other authors declare no conflicts of interest.

## FIGURES, TABLES, and SUPPLEMENTAL MATERIAL

FIG 1, PDF file, 4 MB

FIG 2, PDF file, 2.2 MB

FIG 3, PDF file, 17 MB Fig S1, PDF file, 13.4 MB Fig S2, PDF file, 205 KB

Tables 1-2 and Tables S1-S2 XLSX file, 35 KB

## METHODS

### Study Design

Patient samples were obtained from two academic medical centers with established ILD programs including regular presentation of subjects at an multidisciplinary discussion. Diagnoses for patients with ILD were classified based on multidisciplinary discussion including detailed clinical, radiographic and pathologic information. Biobanked serum was collected and stored by investigators at each site through established research protocols at each center (A.K.S., P.W. at UCSF and A.S., I.N., M.S. at University of Chicago). All research subjects had samples drawn as described under Institutional Review Board (UCSF IRB 10-02467, UCSF IRB 10-01592 & University of Chicago 14163A-AM059).

Two authors (V.U. and C.T.L.) collapsed diagnostic categorization of ILD to shared data categories to facilitate data merging while maintaining the integrity of the original diagnosis. The original center labels and shared diagnostic categories for this manuscript are described in **Supplementary Table 2A**. Control samples were collected from multiple laboratories and represented blood banked donors in San Francisco and New York. Reference samples were collected, aligned, and processed as in (Vazquez et al., 2020, 2022), and serum from these reference samples were re-run with the entire ILD cohort included in this study. A subset of samples of patients with rheumatoid arthritis (RA) without associated ILD were run and additional samples of subjects without known ILD were available from the University of Chicago cohort. Chronic inflammatory disease is established in RA patients from this cohort; the remaining were a combination of relatives of subjects with ILD not known to have ILD themselves (n=16) in whom other diseases were not ruled out and organ donors (n=15). All samples not in the reference group of 138 samples contributed to candidate selection if their Z-scores were above the cutoff. Self-identified race and ethnicity, biological sex, and age ranges for screened subjects are described in **Table 1**. Specific metadata assignments including shared diagnostic label, age, gender, and self-identified race and ethnicity are in **Supplementary Table 2B**.

### PhIP-Seq protocol and Analysis

The PhIP-Seq phage display system was conducted as previously described (Vazquez et al., 2020, 2022). Human serum was incubated with 10^10^ plaque forming units of a phage-display library tiling all open reading frames of the human genome, which was incubated overnight, and precipitated using protein A/G magnetic beads, and used to infected *E. coli* for selective amplification of infective phage and next generation sequencing (Vazquez et al., 2020, 2022). Some samples had sufficient serum available and arrayed for multiple technical replicates; where multiple technical replicates were performed, the maximum PhIP-Seq value collapsed to gene annotations was selected for candidate autoantigen selection. Candidate autoantigens were selected by identifying genes with PhIP-Seq Z scores greater than 17 within the screened cohort and excluding genes in the blood bank control samples (n=138) that had any Z scores greater than 17. Candidates were further filtered as being found 10 times between both cohorts and found at least once in each cohort. Previously described autoantigens were identified completing a literature review and PhIP-Seq data for those candidates was plotted. Resulting Z score data for genes determined to be autoantigens was used to construct two random forest classifiers R version 4.1.1, tidyverse (tidyverse_1.3.1), tidymodels (tidymodels_0.1.4), vip (vip_0.3.2), and pROC (pROC_1.18.0). University of California San Francisco (UCSF) is “Center 1” and University of Chicago is “Center 2” for the purposes of text **Figure 1**. For the inter-center classifier, the blood bank control samples were separated into two groups and paired with one or the other cohort and confidence intervals are displayed from pROC. “Not ILD” was a classification only used at the UCSF center and is visually represented in **Figure 3** but excluded from classifier training given it was not a used category at both centers (**Figure 1**). The “Not ILD” categorization was re-coded to Interstitial Lung Abnormality (ILA) for this manuscript after chart review by A.K.S. Network analysis was conducted using identified candidate antigens as the Z score cutoff for screening candidate antigens and using a Z-score cutoff of 50 for peptide data. A function was created in R to define co-occurrence associations by patient designation for Z score transformed PhIP-Seq data and the resulting plot was created with the Igraph package (igraph_1.3.1).

### Predicted protein structure

Uniprot was utilized to obtain a CDHR5 protein amino acid sequence. Alignments of 11 CDHR5 peptides from **Figure 2D** was completed without incorporating secondary structure information using DECIPHER. Alpha fold (https://alphafold.ebi.ac.uk/) was employed to visualize predicted secondary structure for this protein using the CDHR5 sequence from Uniprot (accession A0A7L0FKT8) (Jumper et al., 2021).

### Validation by RLBA, RNA sequencing, and Immunohistochemistry

Radioligand binding assay (RLBA) was used to validate CDHR5 by translating 35S-labeled CDHR5 using the rabbit reticulocyte lysate system and quantifying the immunoprecipitation with patient serum as described previously (Shum et al., 2013). The CDHR5 vector was prepared by cloning the full length cDNA sequence (XM_011520188) led by the Kozak sequence (GCCACC) into the pTNT vector (L5610, Promega). Commercial anti-CDHR5 (PA5-89483, Invitrogen) was used as the positive control. CDHR5 positivity was defined as 3 standard deviations above the mean of the control subjects as described in the text. A total of 699 samples were tested via RLBA with insufficient serum available from 5 of the original samples screened by PhIP-Seq.

Lung tissue from biopsy specimens for 2 CDHR5 autoreactive subjects were available for sequencing and immunohistochemistry. Samples underwent RNA extraction and were subjected to bulk RNA sequencing as described previously using three age matched unused donor lung tissue as described by (Bhattacharya et al., 2021). These same subjects were used for immunohistochemistry for CDHR5 using a commercially purchased antibody (HPA009081, Sigma Aldrich), and the findings discussed in the text were described by a trained pathologist (K.D.J.).

The authors of this manuscript designed and implemented all aspects of the study, including sample selection, computational methods, data analysis, and manuscript preparation.

### Data Availability

The bulk RNA sequencing for **Figure 3** has been uploaded to the NIH Sequence Read Archive (PRJNA956979). The PhIP-Seq data itself will be made available through public access keys from the Chan Zuckerberg Biohub that can be requested from the DeRisi lab. This data will be available to all public requests at the time of manuscript acceptance and can be made available for reviewers as needed in the interim.

## FIGURES AND LEGENDS

**Supplementary Figure 1:**
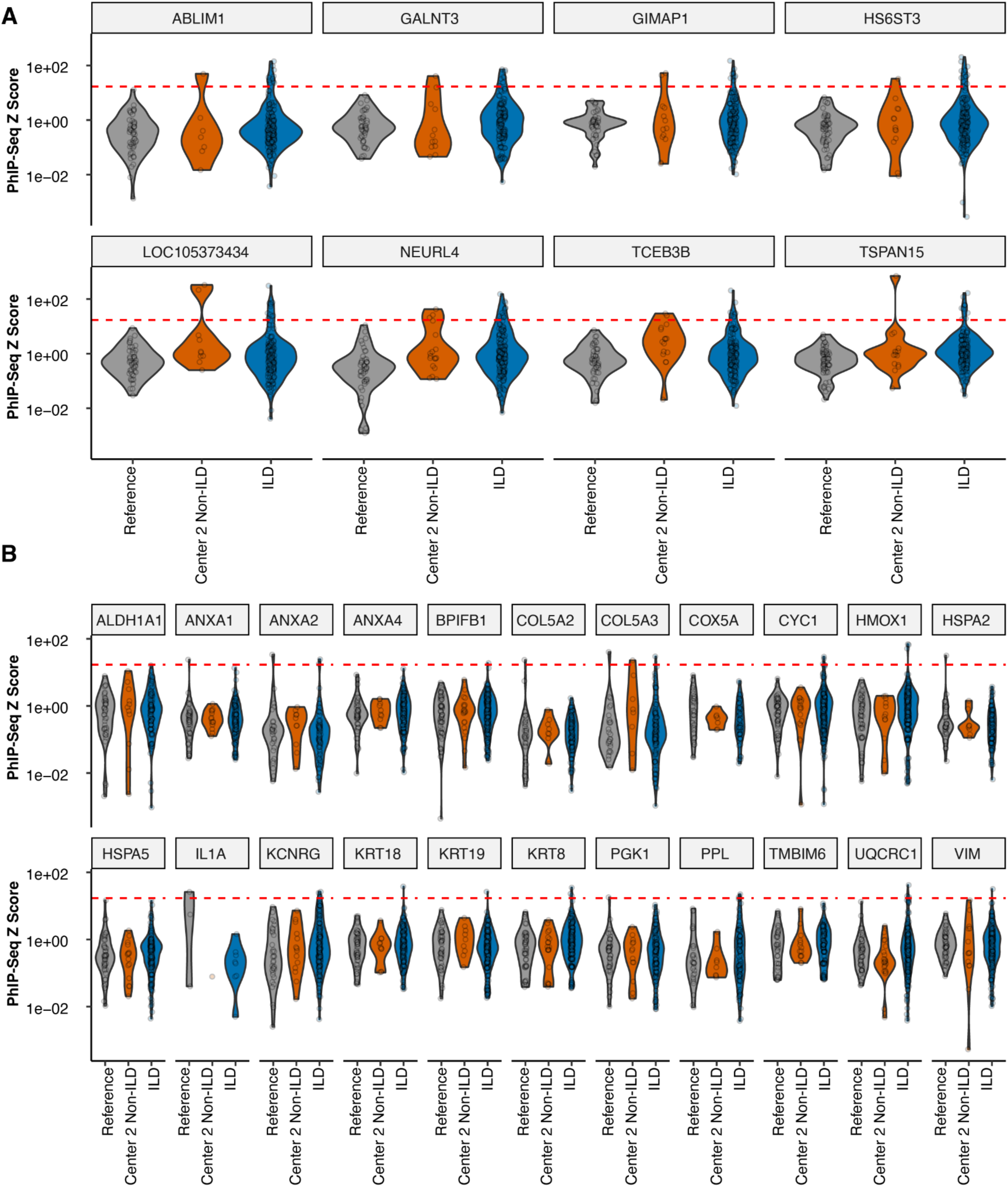
PhIP-Seq assessment of autoantibodies in patients with ILD. **(A)** PhIP-Seq derived autoantibody candidates with high autoreactivity in the Center 2 Non-ILD group. **(B)** Assessment of previously described lung autoantibodies by PhIP-Seq with Reference, Center 2 Non-ILD, and ILD populations as described in **Figure 1**. The red line indicates a Z score cutoff (17) used for candidate picking. n=138 reference, n=398 participants with ILD, n=46 participants in the Center 2 Non-ILD (e.g. 15 patients with rheumatoid arthritis (RA) without known ILD and 31 subjects without ILD from Center 2).

**Supplementary Figure 2:**
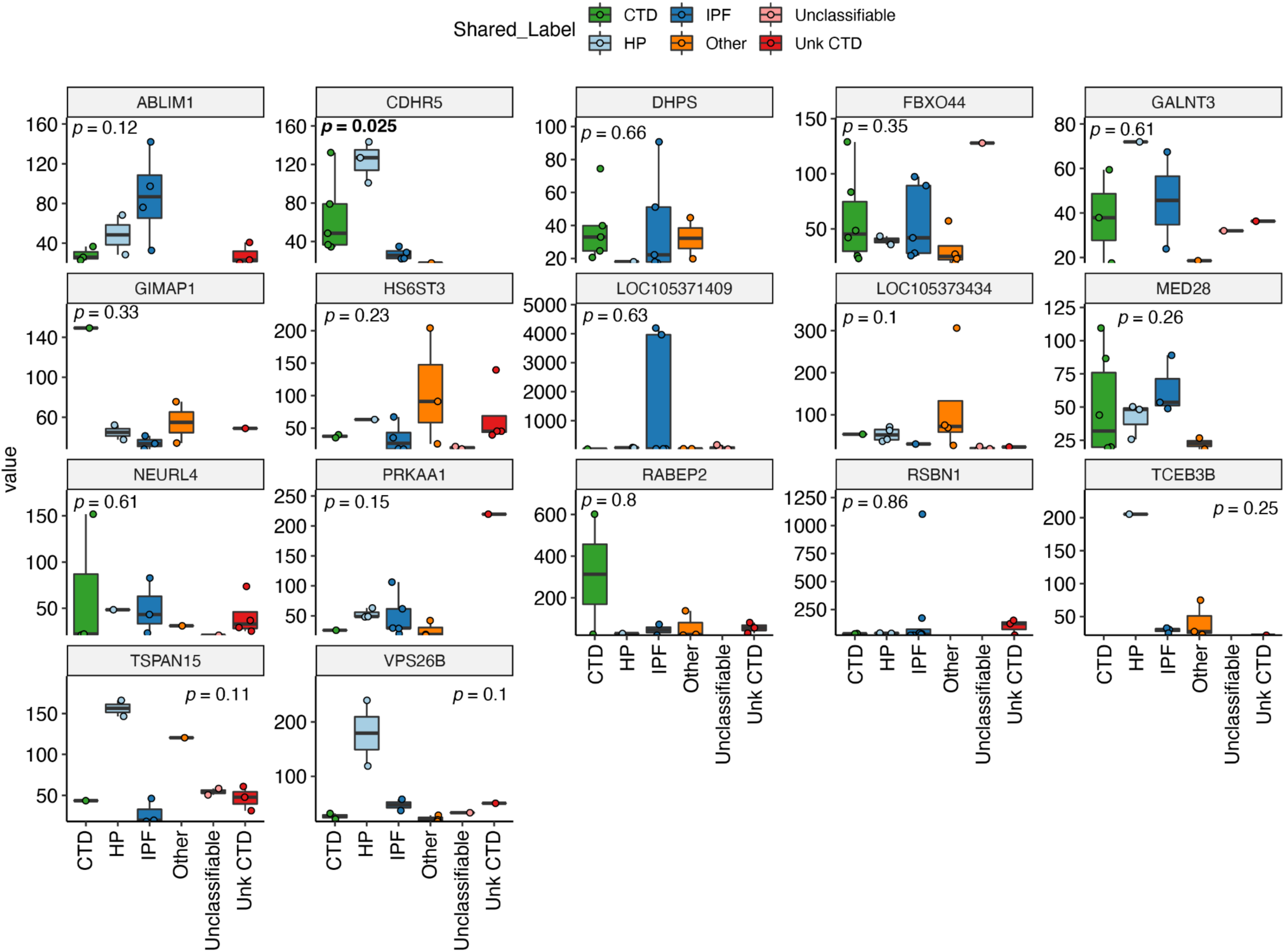
PhIP-Seq positive targets by absolute Z scores by disease categorization. PhIP-Seq positive hits were plotted by absolute value by sample. One-way Kruskal-Wallis ANOVA *p*-value is displayed for values across all candidates comparing the groups on the x-axis to the values for PhIP-Seq Z scores displayed on the y-axis. n=8-18 samples from n=8-13 subjects with ILD for each candidate autoantibody.

